# Tracing Evolutionary Ages of Cancer-Driving Sites by Cancer Somatic Mutations

**DOI:** 10.1101/2020.02.09.940528

**Authors:** Xun Gu, Zhan Zou, Jingwen Yang

## Abstract

Evolutionary understanding of cancer genes may provide insights on the nature and evolution of complex life and the origin of multicellularity. In this study, we focus on the evolutionary ages of cancer-driving sites, and try to explore to what extent the amino acids of cancer-driving sites can be traced back to the most recent common ancestor (MRCA) of the gene. According to gene phylostraigraphy analysis, we use the definition of gene age (tg) by the most ancient phylogenetic position that can be traced back, in most cases based on the large-scale homology search of protein sequences. Our results are shown that the site-age profile of cancer-driving sites of TP53 is correlated with the number of cancer types the somatic mutations may affect. In general, those amino acid sites mutated in most cancer types are much ancient. These sites frequently mutated in cancerous cells are possibly responsible for carcinogenesis; some may be very important for basic growth of single-cell organisms, and others may contribute to complex cell regulation of multicellular organisms. The further cancer genomics analysis also indicates that ages of cancer-driving sites are ancient but may have a broad range in early stages of metazoans.

## Introduction

Cancer occurs in almost all metazoans in which mutated somatic cells proliferate without any control, suggesting that the mechanisms of cancer are deep-rooted in evolutionary history (Weinberg 1983; Nesse and Williams 1998; Merlo, et al. 2006). Hence, evolutionary understanding of cancer genes may provide insights on the nature and evolution of complex life and the origin of multicellularity. Using the phylostratigraphic approach, Domazet-Lozo and Tautz (Domazet-Loso and Tautz 2010) found that evolutionary ages of many cancer genes (gene ages for short) can be traced back to the transition period from unicellular organisms to multicellular metazoans. This so-called cancer-as-atavism hypothesis has been verified by a number of follow-up studies (Davies and Lineweaver 2011; Domazet-Loso, et al. 2014; Chen, et al. 2015; Greaves 2015; Chen, et al. 2018; Trigos, et al. 2018; Makashov, et al. 2019; Trigos, et al. 2019).

As many cancer genes are responsible for the cellular cooperation necessary for multicellularity that also malfunction in cancer cells (Hanahan and Weinberg 2000, 2011), the connection between cancer and the emergence of multicellularity can be revealed through a detailed evolutionary analysis of amino acid sites at which many somatic mutations were detected from multiple cancer samples, i.e., cancer-driving sites. In this study, we focus on the site-age problem, i.e., under a metazoan phylogeny, to trace the (human wide-type) amino acid of a cancer-driving site to a particular ancestral node. Our goal is to explore to what extent the (human) amino acids of cancer-driving sites can be traced back to the most recent common ancestor (MRCA) of the gene. i.e., site ages of cancer-driving sites approach to the gene age.

## Results and Discussion

### Evolutionary age of an amino acid site (site-age)

According to the common practice in gene age (or gene phylostraigraphy) analysis (Domazet-Loso and Tautz 2010), we use the definition of gene age (***t_g_***) by the most ancient phylogenetic position that can be traced back, in most cases based on the large-scale homology search of protein sequences. We extend the concept of gene age to amino acid sites, using the human protein sequence for illustration. Under a species phylogeny that is biologically known or can be reliably inferred from the multiple sequence alignment (MSA), the phylogenetic age of an amino acid site (***t_s_***) is defined as the most ancient node within the lineage that ranges from the phylogeny root to the human, at which the human amino acid type still remains. In other words, ***t_s_*** indicates the most ancient phylogenetic position to which this human-type amino acid residue appeared can be traced back.

Fig. 1 shows the phylogenetic tree inferred from the multi-sequence alignment of TP53 proteins in metazoans, using the unicellular protist *M. brevicollis* as outgroup. According to this phylogeny, the lineage from the human to the root of animals can be divided into several intervals defined by the corresponding internal nodes. With the help of ancestral sequence inference (Yang, et al. 1995; Griffiths and Tavare 2018), it is straightforward to determine the site-age for each amino acid site (see Fig.1 for illustration). For instance, the overall site-age profiles of TP53 are shown in Table 1.

**Figure 1.**
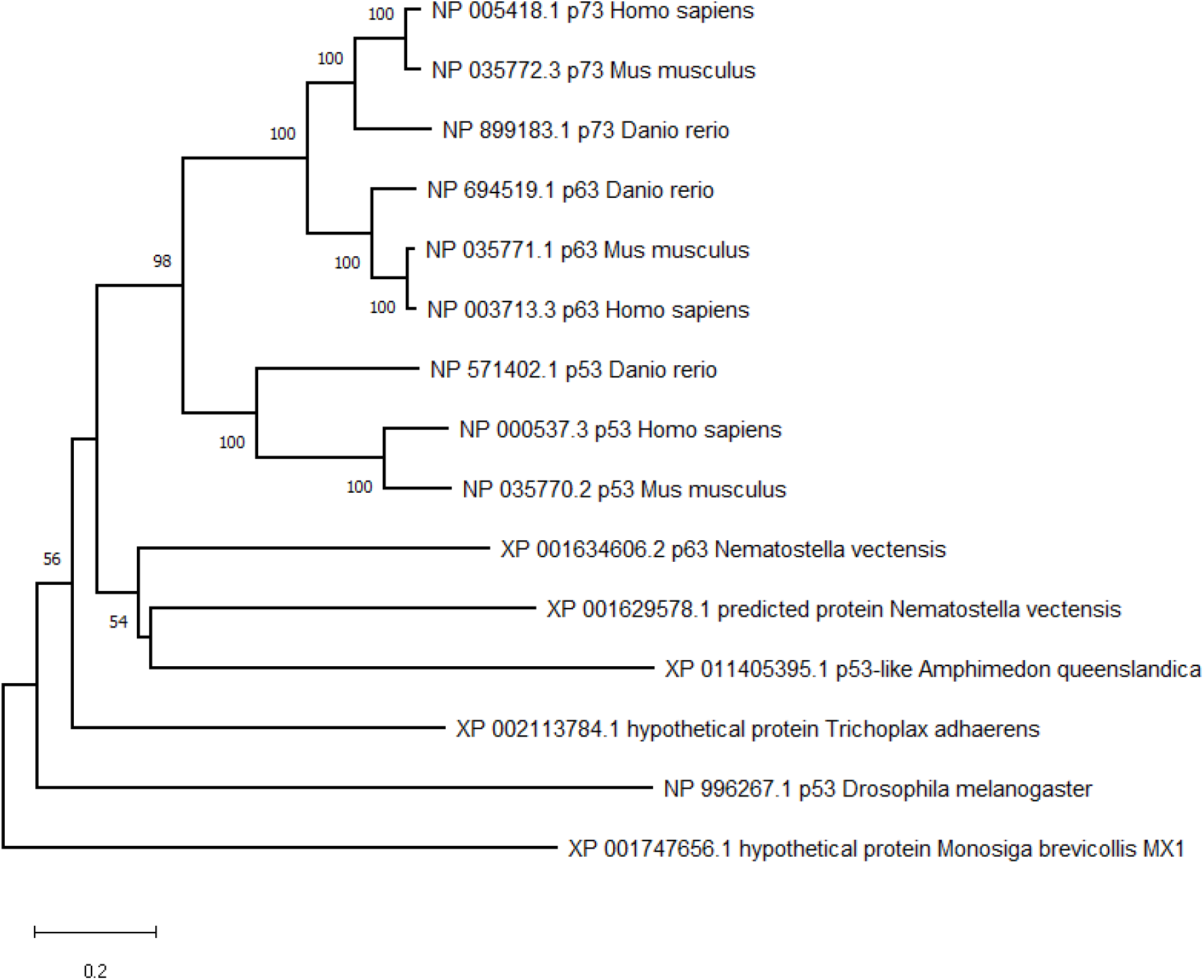
The phylogenetic tree inferred from the multi-sequence alignment of TP53 proteins in metazoans. The phylogenetic tree was inferred using MEGA X with the Neighbor-Joining method. The percentage of replicate trees in which the associated taxa clustered together in the bootstrap test (1000 replicates) are shown next to the branches. The unicellular protist *M. brevicollis* was set as outgroup.

**Table 1.** Site-age profiles of TP53 gene in different phylogenetic groups

| *Datasets* | *Holozoa* | *Bilateria* | *Chordates* | *Teleost* | *Mammals* |
| --- | --- | --- | --- | --- | --- |
| *Amino acid sites* |  |  |  |  |  |
| All sites (393 sites) | 20.0% | 18.5% | 10.0% | 17.8% | 33.7% |
| Kan (57 sites) | 37.3% | 21.6% | 13.7% | 19.6% | 7.8% |
| *Cell types (all 393 amino acids included)* |  |  |  |  |  |
| <20 | 3.5% | 4.6% | 10.3% | 14.7% | 67.0% |
| 20-30 | 15.7% | 11.6% | 14.1% | 25.9% | 32.7% |
| >30 | 39.4% | 24.6% | 16.9% | 12.7% | 6.4% |

### Sites with more cancer somatic mutations tend to have more ancient site-ages: a case study of TP53

Cancer-related amino acid sites, or cancer-driver sites, refer to those who have many recurrent somatic mutations (denoted by *Z*) compiled from different cancer samples. According to the cancer-as-atavism hypothesis, cancer-driver sites tend to have ancient site-age as cancer somatic mutations are functionally harmful, causing cellular genetic instability or deconstruction of cell-cell communications. To test this notion, we use fifty seven amino acid sites of TP53, at which cancer somatic mutations were detected by Kan et al (Kan, et al. 2010) with a strict selection criteria in cancer genomic analysis. Impressively, we observed that the site-ages of cancer-related sites are considerably enriched in the ancient stages of animals such as Holozoa, and Bilateria (Table 1). We used a χ^2^-test and found that the age profile of cancer-related sites differs significantly from the age profile of all sites (P-value<0.001). We thus conclude that these sites may play fundamental roles in backbone cellular genetic program and multicellularity, and somatic mutations occurred at these sites facilitate the cancer progression.

### Ancient TP53 somatic mutation sites tend to affect more cancer types

It is well-known that somatic mutations in TP53, one of most important tumor suppressor genes, have been found in many human tumor types (Greenblatt, et al. 1994). We use UMD TP53 database (http://p53.fr/) that compiles most somatic mutations in TP53 reported in human cancers. While cancer somatic mutations have been found over 90% amino acid sites of TP53, only a small number of sites with somatic mutations from many cancer types. There are two empirical measures: one is the number of case-reports in UMD TP53 database that include somatic mutations occurred at an amino acid site, and the other is the number of cancer types in those case reports. We adopt the second measure in our analysis because it removed the potential redundancy in different case-reports; yet the results are virtually the same (not shown).

Our results are shown in Table 1. It appears that the site-age profile of cancer-related sites of TP53 is correlated with the number of cancer types the somatic mutations may affect. In general, those amino acid sites mutated in most cancer types are much ancient. For instance, most sites with somatic mutations affecting more than 30 cancer types originated from *M. brevicollis*, early metazoan or early vertebrates. These sites frequently mutated in cancerous cells are possibly responsible for carcinogenesis; some may be very important for basic growth of single-cell organisms, and others may contribute to complex cell regulation of multicellular organisms.

### Cancer genomics analysis

To further test the notion that amino acid sites frequently mutated in cancerous cells tend to have more ancient site-ages, we extracted all non-synonymous somatic mutation sites data from COSMIC database (release 63). A total of 17381 sites, each of which has received somatic mutations from at least two samples (z=2 or more), were used in our study, among which 14982 sites are for *z*=2, 2281 for *z*=3, 359 sites for *z*=4, and 821 sites for z=5 or more, respectively. Based on the metazoan phylogeny shown in Fig.1, we inferred the evolutionary ages of all these sites.

As summarized in Table 2, we calculated the site-age profiles for amino acid sites with different counts (*z*) of somatic mutations. For comparisons, we randomly selected 10000 amino acid sites from the human genome to estimate the genome background of site-age profile. It appears that the site-age profile for sites with z=2 is statistically not significant from the background (P-value >0.05). This result is reasonable because most of those sites are passengers so that the corresponding age-site profile is virtually a random sample from the genome background. While the site-age profile for sites with *z*=3 is statistically significant marginally (P-value <0.01), those for sites with *z*=4 and z>=5 are highly statistically significant (P-value <10^-8^). In other words, for those sites with large z counts that are likely to be cancer-driving, their site-ages tend to be more ancient.

**Table 2.** Phylogenetic site-age profiles of amino acid site categories with different counts of cancer somatic mutations (CSM) of somatic mutations.

| **CSM** | **Sites** | *Holozoa* | *Bilateria* | *Vertebrates* | *Tetrapoda* |
| --- | --- | --- | --- | --- | --- |
| *Z=2* | 14982 | 8% | 11% | 27% | 54% |
| *Z=3* | 2281 | 15% | 18% | 28% | 39% |
| *Z=4* | 359 | 25% | 19% | 30% | 26% |
| *Z≥5* | 821 | 28% | 22% | 32% | 18% |
| *Background* | 10,000 | 6% | 9% | 27% | 58% |

We further analyzed a list of 538 known cancer genes. Many well-known cancer genes whose site-ages for sites with z=3 or more can be traced back to the origin of metazoans (the *Holozoa* group); some examples are shown in Table 3. Meanwhile, Table 4 shows some examples that cancer-driving sites can be only tranced back to the emergence of *Bilateria* or early stage of vertebrates. Together, we conclude that ages of cancer-driving sites are ancient but may have a broad range in early stages of metazoans.

**Table 3.**
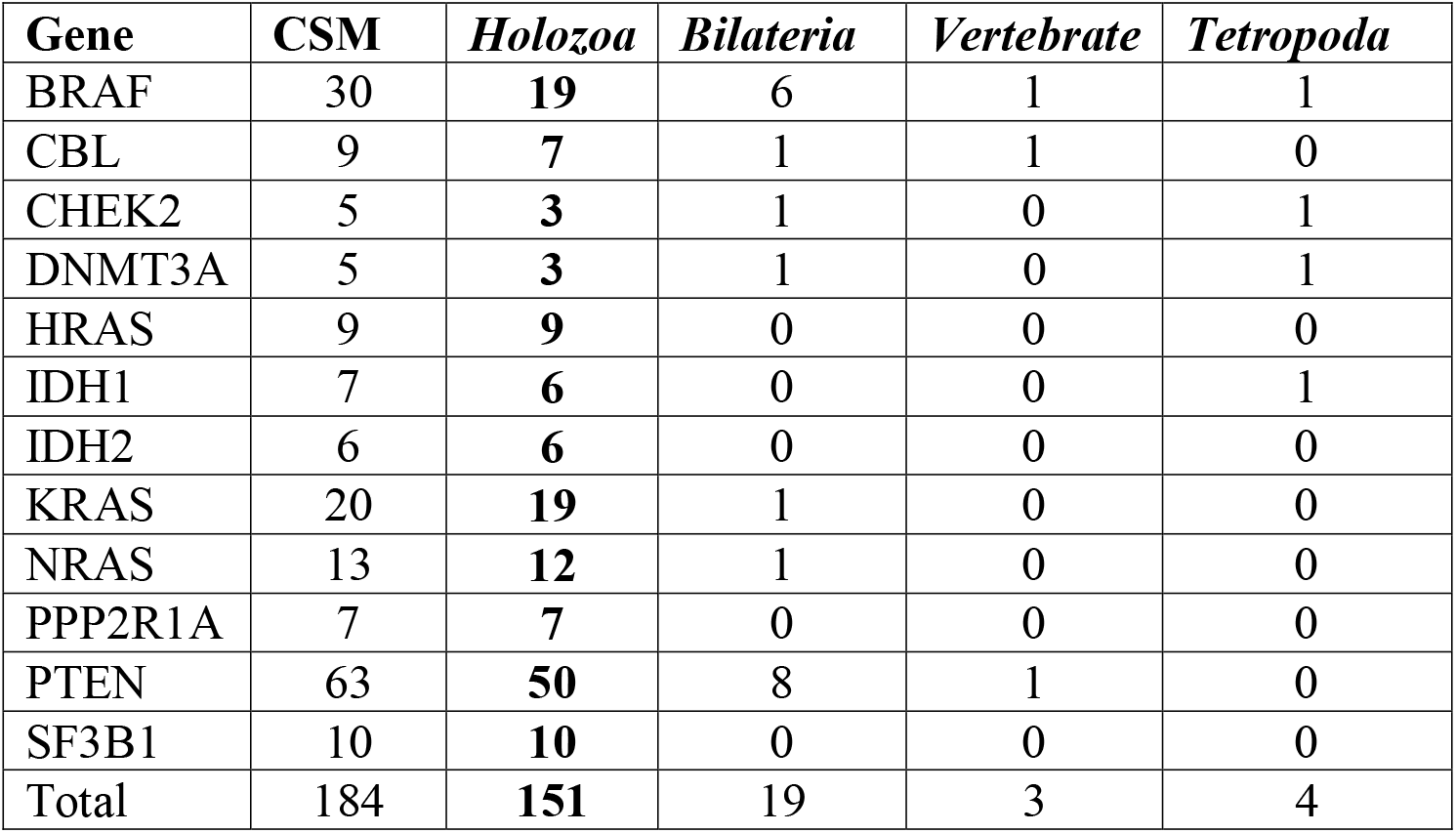
Some examples of cancer genes that can be classified as the Holozoa group

**Table 4.** Some examples of cancer genes that can be classified as the Bilateria group

| **Gene** | **CSM** | ***Holozoa*** | ***Bilateria*** | ***Vertebrate*** | ***Tetrapoda*** |
| --- | --- | --- | --- | --- | --- |
| ABL1 | 14 | 0 | **10** | 4 | 0 |
| CTNNB1 | 34 | 0 | **23** | 10 | 1 |
| EGFR | 60 | 0 | **39** | 16 | 5 |
| EZH2 | 7 | 0 | **6** | 0 | 1 |
| PTPN11 | 13 | 4 | **8** | 1 | 1 |
| SMAD4 | 10 | 0 | **8** | 2 | 0 |
| TSHR | 15 | 0 | **9** | 5 | 1 |
| Total | 153 | 4 | **103** | 38 | 9 |

## Materials and Methods

The protein sequences of most species we used were downloaded from Ensembl (Ensembl 65) and NCBI database. Orthologous genes of these species for human *p53*, *p63* and *p73* genes were searched by BLASTP method. Orthologs of human *p53*, *p63* and *p73* genes in other 11 Drosophila species were obtained by online BLASTP searching in FlyBase website (http://flybase.org). The best hit of each search with *E*-value less 10^-5^ was selected as the homolog of the human *p53*, *p63* and *p73*. than Multiple protein sequence alignment for p53 gene family was generated by MUSLE (Edgar 2004). ClustalW and T-coffee programs were also used to validate the results of multiple sequence alignment (Thompson, et al. 1994; Notredame, et al. 2000). The phylogenic tree for p53 gene family was constructed by neighbor-joining (NJ) method (Saitou and Nei 1987) with Poisson correction distance in MEGA X software (Kumar, et al. 2018). Bootstrap test with 1,000 resampling was used to evaluate the reliability of tree topology for p53 gene family (Felsenstein 1985). We applied the choanoflagellate *Monosiga brevicollis* as outgroup to determine the root of the phylogeny.

Age of each p53 protein site was defined according to ancestral inference of p53 protein sequence by maximum parsimony in MEGA5.0. The evolutionary age was classified into 11 groups, namely Hominidae, Hominoidea, Primate, Mammalia, Tetrapoda, Fish-1, Euteleostomi, Vertebrata, Chordata, Metazoa and Holozoa. Hominidae group means that the amino acid site in p53 originates from Hominidae, while Holozoa represents that the amino acid site of p53 originates from protist *M. brevicollis*, hereinafter and so on. Here the difference between Fish-1 and Euteleostomi groups is that p53 amino acid sites in the former group were earliestly found in zebrafish, while amino acid sites in the latter group were also found in other teleostome species, like Fugu, stickleback and Tetradon. We used age of site conservation to define conservation level of each p53 protein site. Phyletic age of conservation degree was also divided into 11 groups mentioned above. Specifically, p53 amino acid sites in Hominidae group are totally conserved in Hominidae (without any amino acid change), and in Vertebrata group means these sites are completely conserved in vertebrate species, and so on.

TP53 somatic mutation data prevalence in human cancer were downloaded from UMD TP53 database (http://p53.fr/). This resource compiles all TP53 gene variations reported in human cancers with annotations on tumor phenotype, patient characteristics and functional impact of mutations.

The list of 538 known cancer genes was compiled by merging the list of 253 cancer genes with the missense mutation type from the Cancer Gene Census Tier 1 (COSMIC GRCh37, V89)(Tate, et al. 2019), 127 significant mutated genes (SMGs) reported by Kandoth *et al*. (Kandoth, et al. 2013), 260 SMGs reported by Lawrence *et al*. (Lawrence, et al. 2013) and 299 cancer driver genes reported by Bailey *et al*. (Bailey, et al. 2018). We extracted the cancer somatic mutations of 10,224 cancer donors from TCGA PanCanAtlas MC3 project (https://gdc.cancer.gov/about-data/publications/mc3-2017), which include 3,517,790 somatic mutations in coding region and 2,035,693 of which caused amino acid changes in cancers (missense mutations). After filtering some redundancies according to the criterion of Bailey *et al*. (Bailey, et al. 2018), 9,078 samples with 1,461,387 somatic mutations in coding region and 793,577 missense mutations were used in our study.

## References

Bailey MH, Tokheim C, Porta-Pardo E, Sengupta S, Bertrand D, Weerasinghe A, Colaprico A, Wendl MC, Kim J, Reardon B, et al. 2018. Comprehensive Characterization of Cancer Driver Genes and Mutations. Cell 173:371–385 e318.

Chen H, Lin F, Xing K, He X. 2015. The reverse evolution from multicellularity to unicellularity during carcinogenesis. Nat Commun 6:6367.

Chen W, Li Y, Wang Z. 2018. Evolution of oncogenic signatures of mutation hotspots in tyrosine kinases supports the atavistic hypothesis of cancer. Sci Rep 8:8256.

Davies PCW, Lineweaver CH. 2011. Cancer tumors as Metazoa 1.0: tapping genes of ancient ancestors. Physical Biology 8.

Domazet-Loso T, Klimovich A, Anokhin B, Anton-Erxleben F, Hamm MJ, Lange C, Bosch TC. 2014. Naturally occurring tumours in the basal metazoan Hydra. Nat Commun 5:4222.

Domazet-Loso T, Tautz D. 2010. Phylostratigraphic tracking of cancer genes suggests a link to the emergence of multicellularity in metazoa. BMC Biol 8:66.

Edgar RC. 2004. MUSCLE: multiple sequence alignment with high accuracy and high throughput. Nucleic acids research 32:1792–1797.

Felsenstein J. 1985. Confidence Limits on Phylogenies: An Approach Using the Bootstrap. Evolution 39:783–791.

Greaves M. 2015. Evolutionary determinants of cancer. Cancer Discov 5:806–820.

Greenblatt MS, Bennett WP, Hollstein M, Harris CC. 1994. Mutations in the p53 tumor suppressor gene: clues to cancer etiology and molecular pathogenesis. Cancer Res 54:4855–4878.

Griffiths RC, Tavare S. 2018. Ancestral inference from haplotypes and mutations. Theor Popul Biol 122:12–21.

Hanahan D, Weinberg RA. 2000. The Hallmarks of Cancer. Cell 100:57–70.

Hanahan D, Weinberg RA. 2011. Hallmarks of Cancer: The Next Generation. Cell 144:646–674.

Kan Z, Jaiswal BS, Stinson J, Janakiraman V, Bhatt D, Stern HM, Yue P, Haverty PM, Bourgon R, Zheng J, et al. 2010. Diverse somatic mutation patterns and pathway alterations in human cancers. Nature 466:869–873.

Kandoth C, McLellan MD, Vandin F, Ye K, Niu B, Lu C, Xie M, Zhang Q, McMichael JF, Wyczalkowski MA, et al. 2013. Mutational landscape and significance across 12 major cancer types. Nature 502:333–339.

Kumar S, Stecher G, Li M, Knyaz C, Tamura K. 2018. MEGA X: Molecular Evolutionary Genetics Analysis across computing platforms. Mol Biol Evol 35:1547––1549.

Lawrence MS, Stojanov P, Polak P, Kryukov GV, Cibulskis K, Sivachenko A, Carter SL, Stewart C, Mermel CH, Roberts SA, et al. 2013. Mutational heterogeneity in cancer and the search for new cancer-associated genes. Nature 499:214––218.

Makashov AA, Malov SV, Kozlov AP. 2019. Oncogenes, tumor suppressor and differentiation genes represent the oldest human gene classes and evolve concurrently. Sci Rep 9:16410.

Merlo LM, Pepper JW, Reid BJ, Maley CC. 2006. Cancer as an evolutionary and ecological process. Nat. Rev. Cancer 6:924–935.

Nesse RM, Williams GC. 1998. Evolution and the origins of disease. Sci Am 279:86–93.

Notredame C, Higgins DG, Heringa J. 2000. T-coffee: a novel method for fast and accurate multiple sequence alignment. Journal of molecular biology 302:205–217.

Saitou N, Nei M. 1987. The neighbor-joining method: a new method for reconstructing phylogenetic trees. Mol Biol Evol 4:406–425.

Tate JG, Bamford S, Jubb HC, Sondka Z, Beare DM, Bindal N, Boutselakis H, Cole CG, Creatore C, Dawson E, et al. 2019. COSMIC: the Catalogue Of Somatic Mutations In Cancer. Nucleic Acids Res 47:D941–D947.

Thompson JD, Higgins DG, Gibson TJ. 1994. CLUSTAL W: improving the sensitivity of progressive multiple sequence alignment through sequence weighting, position-specific gap penalties and weight matrix choice. Nucleic acids research 22:4673–4680.

Trigos AS, Pearson RB, Papenfuss AT, Goode DL. 2018. How the evolution of multicellularity set the stage for cancer. British journal of cancer 118:145–152.

Trigos AS, Pearson RB, Papenfuss AT, Goode DL. 2019. Somatic mutations in early metazoan genes disrupt regulatory links between unicellular and multicellular genes in cancer. Elife 8.

Weinberg RA. 1983. A molecular basis of cancer. Sci Am 249:126–142.

Yang Z, Kumar S, Nei M. 1995. A new method of inference of ancestral nucleotide and amino acid sequences. Genetics 141:1641–1650.

